# Structural and functional brain changes following four weeks of unimanual motor training: evidence from fMRI-guided diffusion MRI tractography

**DOI:** 10.1101/088328

**Authors:** Lee B Reid, Martin V Sale, Ross Cunnington, Jason B Mattingley, Stephen E Rose

## Abstract

We have reported reliable changes in behaviour, brain structure and function in 24 healthy right-handed adults who practiced a finger-thumb opposition sequence task with their left hand for 10 mins daily, over four weeks. Here we extend these findings by employing diffusion MRI to investigate white-matter changes in the corticospinal tract, basal-ganglia, and connections of the dorsolateral prefrontal cortex. Twenty-three participant datasets were available with pre-training and post-training scans. Task performance improved in all participants (mean: 52.8%, SD: 20.0%; group p<0.01 FWE) and widespread microstructural changes were detected across the motor system of the ‘trained’ hemisphere. Specifically, region-of-interest based analyses of diffusion MRI (n=21) revealed significantly increased fractional anisotropy in the right caudate nucleus (4.9%; p<0.05 FWE), and decreased mean diffusivity in the left nucleus accumbens (-1.3%; p<0.05 FWE). Diffusion MRI tractography (n=22), seeded by sensorimotor cortex fMRI activation, also revealed increased fractional anisotropy in the right corticomotor tract (mean 3.28%; p<0.05 FWE) predominantly reflecting decreased radial diffusivity. These changes were consistent throughout the entire length of the tract. The left corticomotor tract did not show any changes. FA also increased in white matter connections between the right middle frontal gyrus and both right caudate nucleus (17/22 participants; p<0.05 FWE) and right supplementary motor area (18/22 participants; p<0.05 FWE). Equivalent changes in FA were not seen in the left (‘non-trained’) hemisphere. In combination with our functional and structural findings, this study provides detailed, multifocal evidence for widespread neuroplastic changes in the human brain resulting from motor training.

## Introduction

The science of motor rehabilitation for patients with brain injury is hampered by a lack of understanding of how experience changes brain structure and function at microscopic and macroscopic scales. A small number of publications have reported such changes in animal models, cross-sectional comparisons, and/or functional measures of brain change [Chang, 2014]. The majority of such studies have reported results from single imaging modalities (e.g. fMRI) with different study protocols and learning tasks, which can make it difficult to interpret the available research to construct a cohesive picture. In addition, despite decades-old reports of electrical activity inducing myelination *in vitro* [Demerens et al., 1996; Ishibashi et al., 2006], and evidence that myelin is continuously turned over *in vivo* [Savas et al., 2012; Yeung et al., 2014], there have been very few longitudinal studies of white matter (WM) change associated with motor learning in humans. In the few works published, changes in diffusion MRI (dMRI) have predominantly indicated possible changes near the GM/WM interface in visuo-motor areas in participants undergoing visuo-motor skill learning [Scholz et al., 2009] and in the frontal lobe of participants learning a balancing task [Taubert et al., 2010]. Curiously, this latter study reported a reduction, rather than increase, in fractional anisotropy (FA), and a variety of decreases in cortical thickness amongst its results. All of these studies have relied on voxel based morphometry (VBM). Unfortunately, when applied to dMRI, VBM can be greatly influenced by the parameters used, and false positive results can arise through registration biases [Jones et al., 2005; Madhyastha et al., 2014; Thomas et al., 2009]. Although VBM is still considered a valuable family of methods, the voxelwise statistics produced can have low statistical power and, in most instances, leave researchers unable to confidently interpret their results in terms of specific tracts [Bach et al., 2014], which precludes insight into precisely which networks are undergoing change.

Diffusion tractography allows measurement of microstructural properties of specific WM tracts, but is a largely unexplored method in this area, potentially because of several limitations that accompany its most common implementation. First, the method traditionally relies on parcellation-based tract classification, which is likely to include aspects of WM tracts that are functionally irrelevant to the learned task [Reid et al., 2016a]. For example, effects of practicing a hand-based motor task would be ideally investigated in only the part of the corticomotor tract responsible for hand movement, but parcellation-based seeding would typically also include fibres responsible for movement of the face, trunk, and legs. Failure to restrict measures to fibres involved with hand use theoretically weakens the method’s sensitivity, and increases the likelihood that any changes found might reflect other processes, such as developmental maturation [e.g. Baek et al., 2013]. Diffusion metrics from tractography are also typically calculated on a whole-tract-mean basis, opening the possibility of missing changes in a subset of voxels or – if artefactual or due to changes in a crossing fibre – falsely concluding that the entire tract has changed. Finally, neither voxel-based morphometry nor traditional tractography integrate functional information, which limits the ability to infer functional relationships between changes seen, brain function, and skill acquisition.

Previously, we reported a multimodal study of motor learning which used fMRI, TMS, cortical thickness and behaviour (task performance) to assess structural and functional brain changes in 24 healthy adults who practiced a non-visual motor task for ten minutes a day for four weeks [Sale et al.,]. These findings suggested that improvements in task performance were driven, at least in part, by changes within the grey-matter, such as long-term potentiation of synapses. These changes appeared to take place in cortical regions that were already responsible for task performance at baseline, rather than representing gained function in neighbouring areas. The amplitude of TMS-evoked motor evoked potentials reported in this previous work increased significantly following training, which could reflect both grey matter changes (either locally within the stimulated brain region, or in remote but functionally related brain regions), improved white-matter conductivity, or both. Changes in cortical thickness of the dorsolateral prefrontal cortex (dlPFC) were also reported, but neither TMS nor fMRI were optimised for investigating this area.

Here, we report the effects of four-weeks of motor learning on the microstructure of the basal ganglia, corticospinal tract, and related intracortical white matter networks. Unlike previous studies, this experiment was carried out with a pure motor-coordination task, and on a dataset for which a variety of brain changes and performance improvements are known to have occurred [Sale et al.,]. To overcome the aforementioned limitations of traditional dMRI methods, we utilised a recently-validated fMRI-guided diffusion tractography method [Reid et al., 2016b] that allows for sensitive measurement of diffusion metrics in specific functionally-relevant tracts. We also performed a series of subanalyses to confirm results were not a reflection of participant selection, tensor fitting algorithms, voxel exclusion criteria, tractography seeding biases, or variable region-of-interest placement. In combination with our functional and structural findings [Sale et al.,], this study provides compelling evidence that neuroplasticity can be induced throughout the motor system by a relatively focussed form of motor training, and interpreted more robustly when measured with a comprehensive multimodal imaging approach.

## Methods

### Participants and Task

Details of the task and participants have been published elsewhere [Sale et al.,]. In brief, 24 participants were recruited (14 female; aged 28.8 ± 1.5 years; range 18 – 40 years) who were all right-handed (laterality quotient 0.92 ± 0.03; range 0.58 – 1.0) as assessed by the Edinburgh handedness questionnaire. Participants practiced a finger-thumb opposition sequence with their left hand for ten minutes each day, for four weeks. Participants were randomised to practice one of two sequences, and instructed to practice as quickly and accurately as possible but not to watch their hand while doing so. We collected MRI (3T Siemen’s Magnetom Trio) and behavioural data for 23 participants immediately before (*baseline*) and after (*post-training*) this period. Task performance was assessed as the number of correct sequences that could be performed in a 30-second period. This was recorded for each hand by video, outside of the scanner, and quantified offline. Participants gave informed consent and the study was approved by The University of Queensland Human Research Ethics Committee.

### Single Participant T1 Templates

In longitudinal analyses, it is possible for variable regions of interest to induce biases that result in false positive or false negative results. To avoid this, diffusion measures which required atlas-based regions of interest utilised *single participant templates* – T1 images for each participant that were unbiased with respect to time point. The steps used to generate these are described in detail elsewhere [Sale et al.,]. In brief, a participant’s T1 images (MPRAGE, 0.9mm isotropic) at pre-and post-training were N4 corrected, intensity normalised, and registered to one another using half of a symmetrical transform, calculated with ANTS Syn [Avants et al., 2008], after skull stripping. This initial template was converted into a sharper template using the *antsMultivariateTemplateConstruction* script. Segmentation was achieved by using ANTs tools with the ANTs NKI template. If the segmentation was not satisfactory, the brainmask was carefully manually edited and this process repeated. Examples of satisfactory and unsatisfactory segmentations can be found with detailed descriptions of this process elsewhere [Sale et al.,]. Cortical labels were calculated for each participant’s templates using *antsJointLabelFusion* with the ANTS IXI and NKI templates. To identify structures of the basal ganglia, single-participant template T1s were processed with volBrain [Manjon and Coupe, 2016], which has been demonstrated to provide accurate delineation of deep GM structures [Næss-Schmidt et al., 2016].

### Diffusion MRI of the Basal Ganglia

We aimed to determine whether changes in microstructure were apparent in the basal ganglia. At both time points, we acquired a 64-direction high-angular resolution diffusion image sequence with b=3000s/mm^2^, whole brain coverage, and 2.34 x 2.34 x 2.5mm spatial resolution. Our HARDI dMRI pipeline has been published in detail elsewhere [Pannek et al., 2012; Pannek et al., 2014]. In brief, dMRI data underwent extensive preprocessing, and constrained spherical deconvolution was used to estimate fibre orientation distributions. We used MRTrix 3 (https://github.com/MRtrix3/mrtrix3) [Tournier et al., 2012] to calculate tensor metrics across the whole brain.

Registration between T1 and diffusion b0 diffusion images was calculated with FSL’s *epi_reg*, using the previously generated single-participant template white-matter segmentation. By applying the inverse of this transform, labels for the basal ganglia were moved into diffusion space. Mean FA and MD values were then sampled within the labels for the globus pallidus, putamen, caudate nucleus, and nucleus accumbens (NAcc). Although volBrain also provides a label for the amygdala, we considered this area unsuitable for analysis as it is often heavily distorted in dMRI images.

We hypothesised that changes would likely be in the right hemisphere, and in specific ROIs, rather than across the entire basal ganglia. We performed a factorial ANOVA to determine whether changes had occurred differentially in any of these regions. This model was as follows:

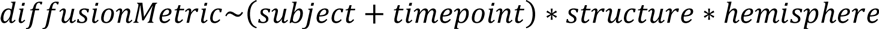

where structure refers to the basal ganglia structure type (putamen, caudate nucleus, etc). This model was used to better account for the non-independence of diffusion metrics between structures in each hemisphere (e.g. left and right putamen) than is possible by treating left and right structures as independent categorical variables. A significant interaction between structure, hemisphere, and time-point was used as the criterion for performing post-hoc paired t-tests to determine which individual ROIs had changed with respect to timepoint. Post-hoc tests were corrected for multiple comparisons using the Holm-Bonferroni method (α < 0.05 FWE). We tested for decreases in MD (one-tailed), but bi-directional changes in FA (i.e. two-tailed), as interpreting FA change is not straightforward in deep cortical grey-matter.

### Diffusion MRI of the Frontal Cortex

Previously, we reported cortical thickness changes in the right (‘trained’) dlPFC in the region of Brodmann’s area 9 [Sale et al.,]. Top-down control of motor output, for error-correction, is predominantly associated with activation of the dlPFC, as well as the caudate nucleus [Chevrier et al., 2007; Kübler et al., 2006]. The dlPFC is thought to connect strongly with premotor areas, including the supplementary motor area (SMA; [Lu et al., 1994]) – a region which displayed changes in fMRI, cortical thickness, and TMS measures after training in the present cohort [Sale et al.,]. For these reasons, we measured FA in connections between the middle frontal lobe and SMA, and the middle frontal lobe and caudate nucleus, with diffusion tractography.

Probabilistic whole-brain tractography was performed on constrained spherical deconvolution images using MRTrix 3.0. For each dataset, 20 million streamlines were generated using anatomically constrained tractography (iFOD2), with *dynamic seeding* and *backtrack* flags, but otherwise default settings. For each hemisphere, we sampled FA for tracts connecting (A) the middle frontal gyrus and caudate nucleus and (B) the middle frontal gyrus and SMA. Labels used were from the single participant T1 templates (see above) to avoid temporal bias. The SMA label included both pre-SMA and SMA-proper. Sampling was weighted according to the output of MRTrix SIFT2 [Smith et al., 2015]. Difference in FA (post-training – pre-training) values violated assumptions of normality and were non-symmetrical. As such, a binomial test was conducted to calculate for an over-representation of FA increases (H_0_: π ≤ 0.5) with Holm-Bonferroni correction for multiple comparisons (α < 0.05 FWE).

### Diffusion MRI of the Corticomotor Tract

We aimed to determine whether diffusion MRI metrics of the right corticomotor tract changed in response to the motor learning task. To focus on functionally-relevant white-matter, we analysed these images with an fMRI-seeded dMRI-tractography protocol that has been published in detail elsewhere [Reid et al., 2016b]. In brief, T1 images were used to generate surface meshes, upon which fMRI activations were calculated (see below). Statistically significant activation within the right S1M1 was used to seed probabilistic tractography from the white matter near the grey-matter/white-matter interface. The same mesh and seed were used for each time point. Tractography was performed on constrained spherical deconvolution images using MRTrix software that had been modified to utilise surface meshes as masks. For each dataset, 40,000 streamlines were generated that passed from the seeding region, through either the posterior limb of the internal capsule or the thalamus (as defined by manually-drawn regions of interest) and terminated in the brainstem. Meshes constrained tractography to the WM. Streamlines were classified as corticomotor or thalamocortical using k-means clustering and streamlines belonging to thalamocortical tracts were discarded. Although complex, this method has recently been shown to reveal more significant and robust relationships between motor performance and diffusion metrics than naïve voxel-based fMRI+dMRI methods, and also provides more coherent tracts than ROI-based classification [Reid et al., 2016b]. Our method here deviated from the previously published version by fMRI task and filtering method (see below). In order to rule out that any changes in the right corticomotor tract were due to biases in registration or tensor-fitting, the procedure was repeated for the left corticomotor tract in all participants who displayed bilateral activation during the functional MRI.

#### Functional MRI

Functional MRI was used to identify functionally-relevant cortical areas for each participant, from which tractography was seeded. We acquired a 2D gradient-echo EPI (TE 28 ms, TR 2670 ms), with online distortion correction and 41 axial slices providing full-brain coverage at a voxel size of 3.28 x 3.28 x 3.3mm (matrix size 64 x 64; FOV 210 x 210 mm; slice gap 0.3mm). Prior to entering the scanner, participants familiarised themselves with the two movement sequences – ‘trained’ and ‘control’ – that they were going to perform within the scanner. Within the scanner, participants performed these finger-thumb opposition movements with their left hand in blocks of 16 seconds, each followed by 16 seconds of rest. During rest blocks, a visual display showed a “Rest” command. At the start of each movement block a visual cue ‐ displaying either the red or blue hand and corresponding movement sequence ‐ notified participants whether they would be performing the trained or the control sequence. This was displayed for 2 seconds, then removed. Participants then performed the required sequence at a rate of two movements per second. As a cue to aid in this timing, a fixation cross flashed at 2 Hz on the screen. A tone also sounded at 2 Hz intervals throughout the acquisition. The last two tones in each movement block were at progressively lower pitches to notify participants that a rest block was imminent. Immediately following completion of each movement block (i.e., following the last fixation cross), a “Stop” command was presented for 1 second, which was then replaced with the “Rest” command, Four consecutive ‘runs’ were performed. Each run consisted of four trained-sequence blocks, four control-sequence blocks, and seven rest blocks. The order of the trained/control sequence blocks was randomised but kept consistent between participants. Correct performance of the sequence was verified by recording the movements with a video camera, which were later reviewed for accuracy.

Each fMRI-session was analysed individually in surface space, using methods and parameters described previously [Reid et al., 2016b]. This included 5mm of surface-smoothing, and motion scrubbing where framewise displacement [Siegel et al., 2014] exceeded 0.9mm. Contrast was set as learned-sequence > rest, for which statistical significance was set at p<0.05 FWE. As this study enrolled healthy adults, we were able to utilise non-linear registration (NiftyReg [Modat et al., 2010]) to propagate labels from the Harvard-Oxford atlas to the mesh in order to crop significant activation to the right S1M1. Manual filtering of significant activation was then carried out, as described previously [Reid et al., 2016b], to retain only the largest ROI near the hand-knob on the right S1M1. For each participant, regions of significant activation from each time-point were combined with logical ‘OR’ to provide an identical seeding region for each time-point’s tractography. This seeding region was expanded such that total seeding area was at least 200mm^2^ in participants who displayed significant fMRI activation, but whose total seeding area was below this size.

#### Statistical Analysis of Tractography

We sought to determine whether FA had changed over time for the corticomotor tracts. Diffusion metrics were sampled for each tract, and the mean taken. Paired t-tests were applied to pre-and post-training tensor values to determine whether change in FA or MD occurred with training. It has become commonplace in VBM analyses to exclude voxels with FA values < 0.2, in order to exclude GM voxels [Bach et al., 2014]. Tractography naturally avoids such voxels, but to maintain comparability with previous works in this area, which have relied on VBM analyses, we repeated our analysis excluding all samples from voxels with an FA < 0.2.

Changes in diffusion metrics were expected to be subtle. Although probabilistic tractography is robust to minor deviations in seeding position (especially with multiple inclusion ROIs), the possibility remains that deviations in tractography and/or errors in tensor fitting could cause a small number of voxels to artificially raise the mean FA of a tract, particularly near the seed point. The nature of axonal ‘all-or-nothing’ signalling and relationships between activity and subsequent myelination [Demerens et al., 1996; Ishibashi et al., 2006] imply that genuine changes in white matter should occur throughout the entire length of a tract. To confirm that any detected changes reflected genuine change throughout the length of the corticomotor tract, each streamline was binned into seven equal-length sections, using a method previously described [Sahama et al., 2015], and intensities were sampled for each section. An example of this binning is shown in Figure 2. Means for each bin were entered into a factorial ANOVA, with *time-point*, *participant*, and *bin* as factors, as well as *participant * time-point and time-point * bin* interactions, using R [R Core Team, 2015]. *Bin* was treated as a categorical, rather than continuous, variable due to the non-linear nature of FA values across the tract length. The significance of the *time-point * bin* interaction in this model was inspected to determine whether any mean change was due to differences in a small cluster of voxels, or local variations in tractography, rather than biologically plausible changes. Criteria for further investigation into any such issues was set as (A) a loss of significance (p≥0.05) for the ‘time-point’ factor, and/or (B) trends towards significance (p<0.1) for this interaction term.

### TBSS

To facilitate comparison with previous works, we also ran tract based spatial statistics on our data. As voxelwise approaches are largely outside of the intended goals of this work, they are detailed and discussed in Supplementary Materials.

## Results

Results are summarised in Table 1. One participant displayed MRI artefacts on the T1 images in the right sensorimotor cortex, and so was excluded from all results reported here, leaving 22 participants. For these 22 participants, increased execution speeds were observed in the post-training assessment for both the trained (Left hand: 53 ± 4.0% [Mean ± SEM]; Right hand: 23 ± 4.2%) and control sequences (Left hand: 7.6 ± 3.0%; Right hand: 17 ± 3.9%). These mean increases were all statistically significant (One-tailed paired T-Test; Holm-Bonferroni corrected p<0.01). The degree of performance improvement was greater for trained than untrained sequence in the left hand (p=1.2E-9) but not in the right hand (p=0.07).

**Table 1.**
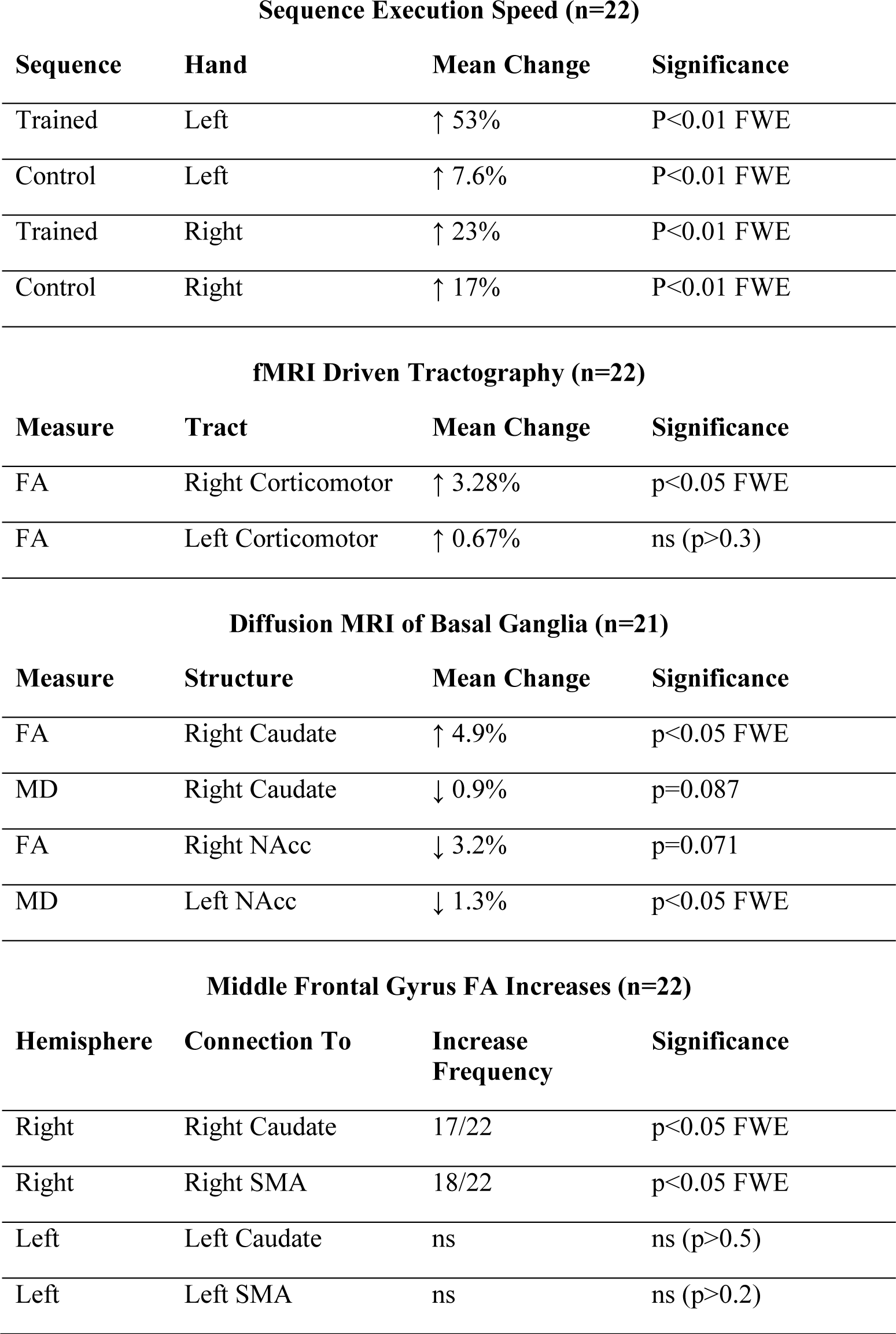
Summary of major findings. Values are rounded to 2 significant figures. Significance is uncorrected for multiple comparisons, except where ‘FWE’ is signified. Abbreviations: FA, fractional anisotropy; FWE, Family-wise error (multiple comparisons) corrected; n, number of participants; ns, not significant; MD, mean diffusivity; NAcc, Nucleus Accumbens; RD, radial diffusivity; SMA, supplementary motor area.

**Sequence Execution Speed (n=22)**
| Sequence | Hand | Mean Change | Significance |
| --- | --- | --- | --- |
| Trained | Left | ↑ 53% | P<0.01 FWE |
| Control | Left | ↑ 7.6% | P<0.01 FWE |
| Trained | Right | ↑ 23% | P<0.01 FWE |
| Control | Right | ↑ 17% | P<0.01 FWE |

**fMRI Driven Tractography (n=22)**
| Measure | Tract | Mean Change | Significance |
| --- | --- | --- | --- |
| FA | Right Corticomotor | ↑ 3.28% | p<0.05 FWE |
| FA | Left Corticomotor | ↑ 0.67% | ns (p>0.3) |

**Diffusion MRI of Basal Ganglia (n=21)**
| Measure | Structure | Mean Change | Significance |
| --- | --- | --- | --- |
| FA | Right Caudate | ↑ 4.9% | p<0.05 FWE |
| MD | Right Caudate | ↓ 0.9% | p=0.087 |
| FA | Right NAcc | ↓ 3.2% | p=0.071 |
| MD | Left NAcc | ↓ 1.3% | p<0.05 FWE |

Table 1. Summary of major findings.
| Hemisphere | Connection To | Increase Frequency | Significance |
| --- | --- | --- | --- |
| Right | Right Caudate | 17/22 | p<0.05 FWE |
| Right | Right SMA | 18/22 | p<0.05 FWE |
| Left | Left Caudate | ns | ns (p>0.5) |
| Left | Left SMA | ns | ns (p>0.2) |

## Diffusion MRI of the Basal Ganglia

No additional datasets were excluded from the diffusion MRI analysis of the basal ganglia (n=22). An example of ROI placement is shown in Figure 1. A significant interaction between structure, hemisphere, and time-point was found (p=0.025) and so post-hoc tests were performed. In the right caudate nucleus, FA increased significantly (4.9%; p=0.002; p-corrected<0.05), and there was a trend towards an MD decrease (-0.9%; p=0.087; p-corrected=ns). In the NAcc, a trend toward a decrease in FA was apparent in the right hemispheric nucleus (-3.2%; p=0.07; p-corrected=ns), and a decrease was seen in MD of the left hemispheric nucleus (-1.3%; p=0.005; p-corrected<0.05). We did not see changes in the globus pallidus or putamen. To test the robustness of these findings, we reran analyses using ANTs-generated basal ganglia labels, instead of those from volBrain, and utilised tensor maps calculated with an older version of MRTrix (2.9), which uses a different tensor fitting method. This reanalysis found a similar pattern of changes (data not shown).

**Figure 1.**
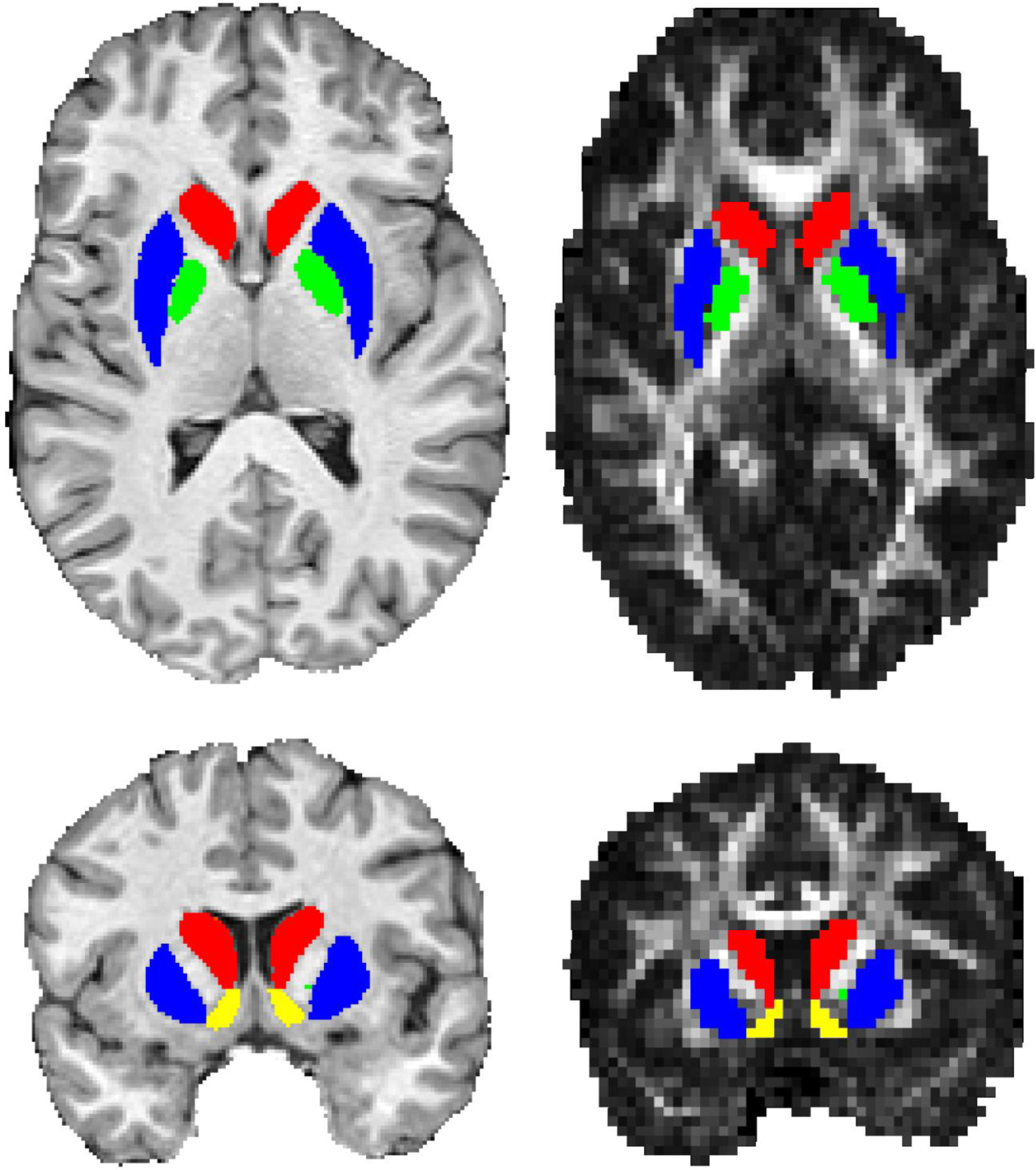
Example of volbrain ROI placement on a single participant template (left) and to a diffusion FA image (right). Colours: Red, Caudate Nucleus; Blue, Putamen; Green, Globus Pallidus; Yellow, Nucleus Accumbens.

## Diffusion MRI of the Frontal Cortex

Wholebrain tractography and related procedures were successful for all available participants. FA increased for tracts connecting the right middle frontal gyrus to (A) the right caudate nucleus in 17 of 22 participants (p < 0.05 FWE; median 1.78% increase), and (B) the right SMA in 18 of 22 participants (p<0.05 FWE; median 1.71% increase). These changes appeared to be driven mainly by numerical decreases in radial diffusivity (median change: Caudate, −0.86%, SMA, −0.99%), rather than increases in axial diffusivity (Caudate, 0.59%, SMA, 0.35%). No significant changes were found in the left hemisphere.

## Diffusion MRI of the Corticomotor Tract

### fMRI

In this study, fMRI was used to delineate hand knob cortical regions for seeding tractography rather than to provide a measure of cortical plasticity. Changes in fMRI patterns with training have been reported in a previous paper [Sale et al.,]. For this reason, the fMRI activation maps for both baseline and post-training sessions were combined for each participant to delineate the hand knob region. Statistically significant S1M1 activation was unilateral in 5 participants and bilateral in the remaining 17 participants. An fMRI activation pattern from a typical participant is shown in Figure 2. All participants had significant S1M1 activation in the right hemisphere; this was used to seed tractography which delineated the right corticospinal tract. For those participants with bilateral activation, the motor region with significant BOLD response in the left hemisphere was used to delineate the left corticospinal tract. Significant activation was also commonly found in the supplementary motor area (21 participants) and superior parietal lobule (20 participants), but such activation was not used for seeding tractography.

### Whole-Tract FA and MD

After training, mean FA of the right corticomotor tract had increased by 3.28% compared with the baseline measurement (p<0.05 FWE; median 2.01%), driven predominantly by a mean 2.2% decrease in radial diffusivity (median 1.34% decrease). MD of the right corticomotor tract was unchanged (p>0.1). FA and MD of the left corticomotor tract were also unchanged (both p>0.05). When the five participants who did not display left-hemispheric fMRI activation were also excluded from the right-hemispheric results, mean change in FA in the right corticomotor tract strengthened to a 3.95% increase (p<0.004 uncorrected; median 3.45%). When these analyses were repeated with a voxel inclusion criteria of FA ≥ 0.2, or with tensor maps calculated with older version of MRTrix (2.9) which uses a different tensor-fitting algorithm, the same pattern of results was obtained (data not shown). Although the seeds were identical for each time point, we sought to alleviate concerns that results were due to some unforeseen complication with fMRI seeding, and reran the analysis defining each corticospinal tract in a much more simplistic manner – tracks connecting the precentral gyrus to the brainstem. A similar pattern of results was seen, albeit with numerically smaller degrees of change in the right hemisphere (data not shown), as one may expect from a substantially less anatomically-specific method [Reid et al., 2016b].

### Along-Tract Analysis

On the basis of an apparent increase in FA, the right corticomotor tract was split into seven equal-length sections from which FA was resampled (Figure 2). A factorial ANOVA fed mean bin-FA values yielded significant effects of *time-point* (p<0.001), and all controlled-for factors (see methods; all p<0.001) but the *bin * time-point interaction* was not significant (p=0.42), implying change was relatively consistent (did not significantly differ) throughout the length of the tract. Unadjusted mean FA increased by 2.10 – 4.3% across the seven bins (Figure 3). This same pattern of results was found when the requirement for voxels to have an FA ≥ 0.2 was applied, or MRTrix 2.9 was used for tensor fitting (data not shown).

**Figure 2.**
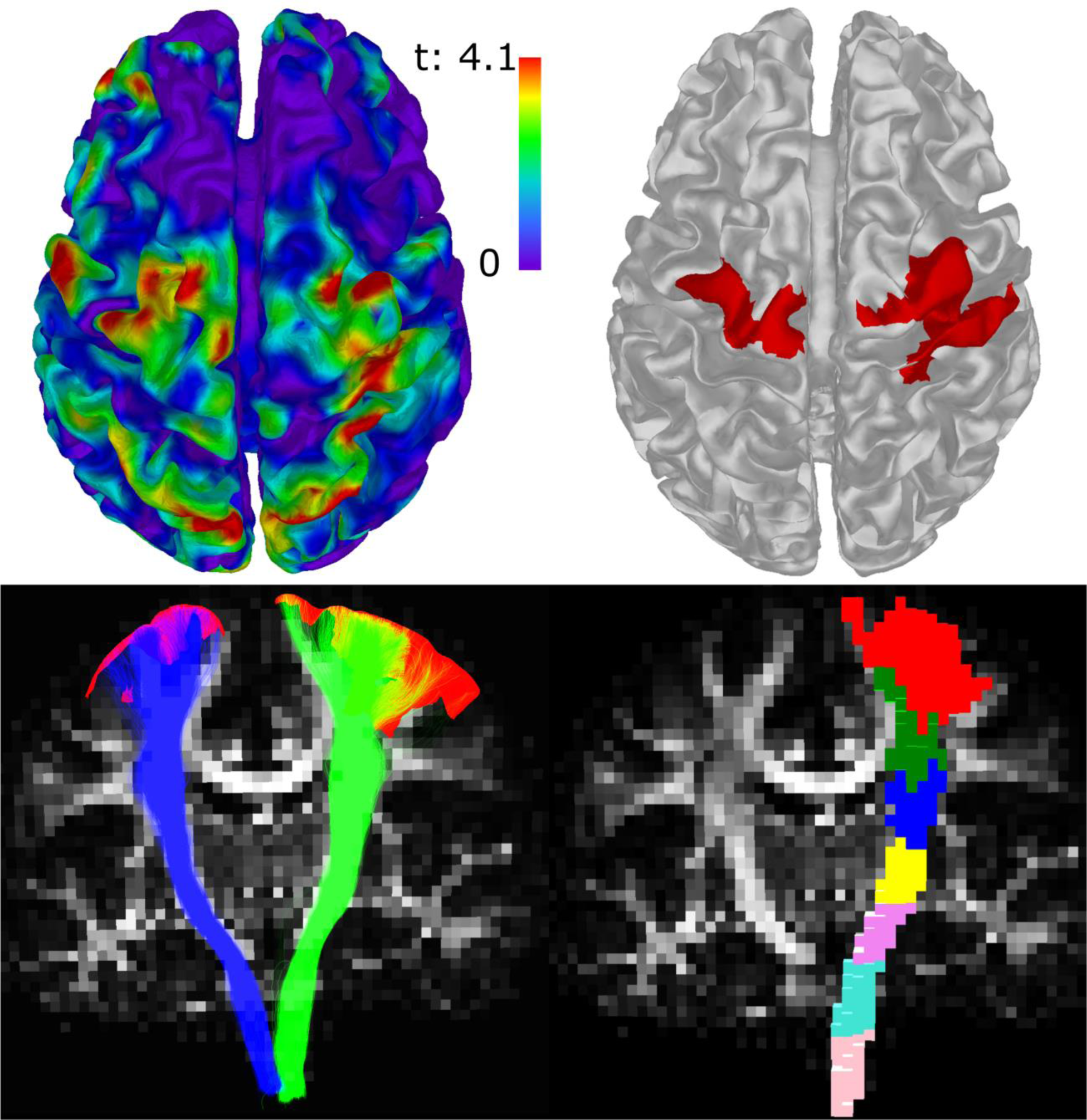
fMRI-tractography for a representative participant. **Top Left:** fMRI t-value map for learned-task versus rest. **Top Right:** Areas of significant fMRI activation (red) after filtering and transforming to the grey-matter white-matter interface (silver; see text). **Bottom Left:** Left (blue) and right (green) corticomotor tracts, eludidated with tractography seeded from the thresholded fMRI (red), overlaid on the fractional anisotropy image. **Bottom Right:** 3D rendering of voxels crossed by the right corticomotor tract, broken into seven bins.

**Figure 3.**
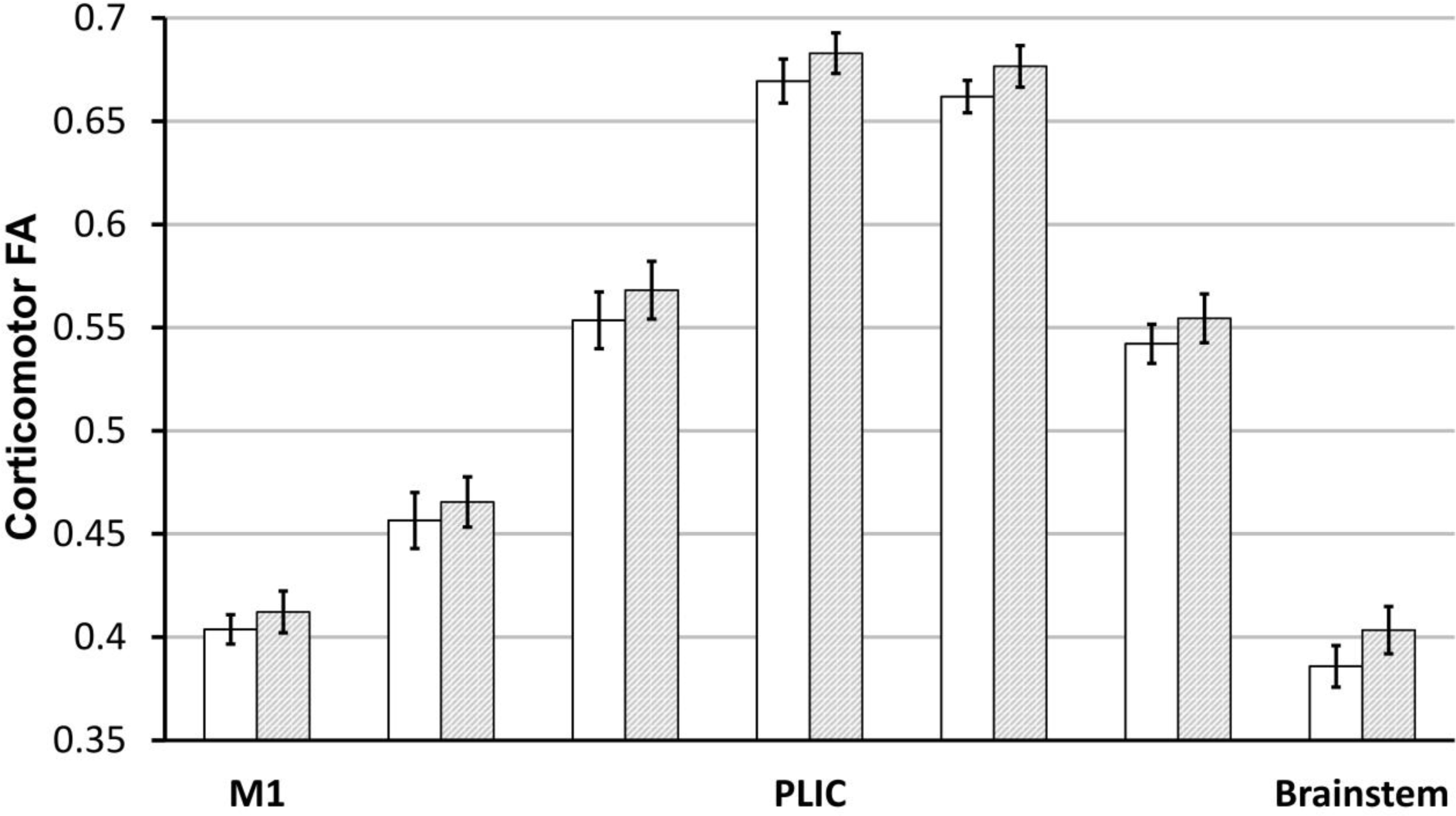
Change in unadjusted FA in the right corticomotor tract, by bin. White and grey bars indicate FA before and after training, respectively. The leftmost bars represent the most superior bin sampled (motor cortex); the rightmost bars represent the most inferior bin sampled (brainstem). Error bars denote SEM. Differences between time points were statistically significant and did not differ by bin. Abbreviations: PLIC, posterior limb of the internal capsule; M1 primary motor cortex.

## Discussion

In this study, 23 right-handed participants practiced a finger-thumb opposition task with their left hand over a four-week period. Neuroimaging revealed a variety of brain changes throughout motor-related areas in the right hemisphere. Previously, we reported reduced fMRI activation, increased TMS motor evoked potentials of the corticospinal tract, and suggested cortical thickness increases throughout the motor system, including the dlPFC. Here we found that FA increased significantly throughout the length of the right corticomotor tract (Figure 3). Investigation of the basal ganglia revealed a significant increase in FA (4.9%) and a trend toward decreased MD (−0.9%) of the right caudate nucleus, as well as significantly decreased MD in the left NAcc (-1.3%) and a trend toward decreased FA of the right NAcc (-3.2%). On the basis of these changes we investigated white-matter connections between the middle frontal gyrus (where cortical thickness changes were reported) and the caudate nucleus (Right hemisphere: FA increase in 77% of participants; Left hemisphere, not significant), and also with the SMA (Right hemisphere: FA increase in 82% of participants; Left hemisphere, not significant). Notably, changes found in all measures were unilateral – appearing solely in the right (‘trained’) hemisphere, with the exception of FA in the nucleus accumbens, which trended towards bilateral changes. Combined with our findings of from TMS, cortical thickness and fMRI changes, the present work provides compelling evidence that neuroplasticity associated with a specific motor learning task occurs throughout the motor system, and can be robustly measured using a multimodal approach.

### Sensorimotor Cortex

Motor output from the brain predominantly originates from the sensorimotor cortex [Borich et al., 2015; Matyas et al., 2010], and it was from the corticomotor tract that we saw an increase in FA, driven predominantly by changes in radial diffusivity. From the same dataset we previously reported changes in cortical thickness, fMRI maps, and TMS maps [Sale et al.,]. The locations of such changes are consistent with brain stimulation work in which areas responsible for motor output of the fingers have been mapped [Penfield and Boldrey, 1937].

Although it is difficult to ascertain precisely what our previously reported changes in cortical thickness represent at a cellular level [Zatorre et al., 2012], they lend credence to the diffusion changes found, for which we can hypothesise a biological origin. The task practiced by our participants involved sustained, rapid, repetitive use of the digits of the non-dominant hand in a manner different from those expected for most daily activities. Given that myelination has previously been demonstrated to occur in response to electrical activity [Demerens et al., 1996; Ishibashi et al., 2006], and we can expect action potentials to traverse the entire length of the corticomotor tract, it is not unreasonable [Zatorre et al., 2012] to surmise that an approximately uniform increase in myelination could result from extensive practice of this task. Although we present this hypothesis tentatively, it is consistent with our observations of altered radial diffusivity, an approximately-uniform change in FA across the tract length, as well as with the TMS measures we previously reported in these participants [Sale et al.,]. Once the technique has sufficiently matured [Alonso-Ortiz et al., 2015], myelin water imaging may provide a useful tool to confirm or refute plasticity of this kind in motor skill learning paradigms. Because of its role, it is likely that the FA changes we have seen in the corticospinal tract reflect the extensive practice of fine finger movements in general, and do not reflect changes arising from this particular sequence.

This study is the first to report tract-specific WM changes in response to motor skill learning, and the first study of its kind in which dMRI changes are supported by concurrent TMS changes. While we are cognisant of the work by Palmer et al [Palmer et al., 2013], who performed another longitudinal study involving fMRI guided tractography, this work focussed on simple strength training rather than motor skill learning, and utilised an alternative approach to tractography seeding and categorisation that is possibly susceptible to cross-sulcal smoothing and other issues [Reid et al., 2016b]. Unlike the present study [Sale et al.,], changes in fMRI activation in the strength training study of Palmer et al. were also absent, as were changes in FA, which one might expect to be more sensitive than changes in MD for much of the corticospinal tract, given its large size and unbranching nature.

### Dorsolateral Prefrontal Cortex and Anterior Striatum

The pre-and post-central gyri are known for their roles in subconscious motor output. By contrast, top-down control of motor output, for error-correction, is predominantly associated with activation of the dlPFC, in which we previously reported cortical thickness changes, and the caudate nucleus [Chevrier et al., 2007; Kübler et al., 2006], in which we saw changes in microstructure. Functional MRI studies have also previously implicated the dlPFC and striatum as playing a key role in motor learning: fast learning of sequential motor tasks modulates activity in these areas [Dayan and Cohen, 2011]. Inhibiting NMDA receptor currents in the striatum has also been shown to severely disrupt motor learning in mice, without disrupting pre-existing motor behaviours or other forms of learning [Dang et al., 2006]. There is also a growing body of evidence that the NAcc plays a role in goal-directed behaviours [Mannella et al., 2013].

Anatomically, both the caudate nucleus and nucleus accumbens receive direct input from the dlPFC, suggesting that changes in the basal ganglia may have been driven by extensive utilisation of top-down motor-control networks, as may be expected when learning a task that requires a high degree of coordination. This hypothesis is supported by our finding of increased FA in connections between the middle frontal gyrus and both the caudate nucleus and SMA, as may be expected from extensive excitation of these pathways. It is also supported by the fact that we did not see changes in the putamen, the other primary input area to the basal ganglia, whose input predominantly originates from more posterior motor areas that play a lower-order role in motor processes [Chevrier et al., 2007].

### Generalisation to the Untrained Hand

Improvements in task performance were seen for both sequences in both hands, despite only one sequence with the non-dominant hand being trained. Such cross-hemisphere generalisation is at odds with one smaller (n=6) previous study employing a similar learning task [Karni et al., 1995], but consistent with findings from several other motor studies utilising other tasks [Hicks et al., 1983; Teixeira, 2000]. Our fMRI analyses revealed that, after training, execution of the trained sequence with the left hand elicited reduced activation relative to the control sequence, *bilaterally* [Sale et al.,]. However, no changes in the left hemisphere were revealed by either structural or diffusion MRI. This discrepancy suggests that altered fMRI and behavioural measures of the ‘untrained’ side may reflect changes to the grey matter, such as LTP or altered neurite density, that were too subtle for changes in either measure. It may also reflect increased intercortical inhibition originating from the right hemisphere, though this seems at odds with the diminished relative activity reported in the right hemisphere. Techniques such as neurite orientation dispersion and density imaging [Zhang et al., 2012], bilateral TMS, EEG, and fMRI tasks targeting the opposite hand may allow future techniques to shine light on this process.

### Strengths and Limitations

Discovering changes in brain function and structure in response to learning has captured the imagination of researchers and the public alike [Malcom, 2015; Schulder and Grabow, 2011; Storr, 2015]. The changes that we have indexed here are relatively subtle. This should be expected – a healthy person performs hundreds of learned complex tasks on a daily basis; the addition of a novel task may be expected to invoke changes that allow, or are represented by, circuit adaptation, but should not be expected to evoke large or permanent changes in brain structure. As expected, practice-induced changes in any such study are subtle, and false positive results are possible if caution is not taken in the analysis and interpretation of data [Thomas et al., 2009].

One strength of the current study was overcoming and controlling for potential sources of error. First, tractography-based studies could conceivably be biased by inconsistent seeding (i.e. inconsistent distance from white/grey matter interface due to registration errors), or changes in diffusion metrics (due to genuine plasticity or image artefacts) of voxels containing grey-matter near seeding locations or in the midbrain. To eliminate the possibility that mean change in FA was due to a subset of voxels shifting the whole-tract mean, we binned the tract into sections and demonstrated that change did not significantly differ at any one location in the tract. This method also ascertained that changes in FA did not simply index change in other fibres which cross the corticomotor tract near the cortical surface. Second, we controlled for variable diffusion tensor fitting by demonstrating a lack of change in corticospinal tracts in the untrained hemisphere, and replicating results using FA images derived with an alternative tensor-fitting algorithm. Finally, and critically, the changes reported were consistent with changes in TMS measures, and backed by increases in skill performance, cortical thickness changes, and altered fMRI activation in the sensorimotor cortex and superior parietal lobule [Sale et al.,].

It is reasonable to assert that the two primary sources of error for ROI-based dMRI analyses are variability in the tensor fitting algorithm and variability in ROI placement. As ROIs were generated on single-participant templates, and propagated back to each time point, it is unlikely that a bias in their placement took place. We also tested an alternative package to generate our ROIs and found similar results. Similarly, utilising a less sophisticated, but more established, tensor fitting algorithm again yielded the same pattern of results.

The pattern of subtle brain changes seen, taken in context of our functional findings [Sale et al.,], suggests that motor learning involves structural brain changes. It should be noted, however, that our study lacked a second follow-up time point, and so it is ambiguous as to whether our detected changes reflect semi-permanent changes that accompany ability improvement, or whether they are a side effect of microstructural (e.g. vascular, dendritic) change that subsides once performance improvement reaches a plateau.

## Conclusion

In response to unilateral motor-coordination training, diffusion MRI revealed changes in the anterior striatum and white matter connections between this region and the right (‘trained’) middle frontal gyrus. White matter connectivity between the right frontal gyrus and right SMA also appeared to improve. Finally, FA increased throughout the length of the right corticospinal tract. These changes were consistent with concurrently recorded changes in fMRI, TMS, cortical thickness, and performance improvements outlined in detail in a separate paper [Sale et al.,]. These results imply that, while some areas may be critical for driving motor learning, this process evokes widespread functional, structural, and microstructural changes across the motor system, and that effective motor rehabilitation schemes may be able to achieve widespread brain changes without necessarily needing to target specific aspects of motor processing individually.

## Acknowledgements

This work was supported by a Training Fellowship from the National Health and Medical Research Council of Australia (APP1012153) awarded to MVS. JBM was supported by an Australian Research Council Laureate Fellowship (FL110100103). The funders had no role in study design, data collection and analysis, decision to publish or preparation of the manuscript. The content is solely the responsibility of the authors and does not necessarily represent the official views of the funding bodies. The authors declare no competing financial interests. We would like to thank Dr. Amanda Robinson, Dr. David Lloyd, and Dr. Daniel Stjepanovic for technical assistance.

## Tract Based Spatial Statistics

### Method

Preprocessed diffusion MRI data (see Main Text) were utilised from 22 participants. Fractional anisotropy (FA) and mean diffusivity (MD) maps were calculated using MRTrix 2.9. A whole-brain analysis using the standard tract-based spatial statistics (TBSS) pipeline [Smith et al., 2006] was conducted. A paired analysis was achieved by subtracting the post-training timepoint from the baseline time-point after registration to the template, and conducting a one-sample T-test with threshold-free cluster enhancement using FSL ‘randomise’ [Smith and Nichols, 2009]. An exploratory re-analysis using a recently introduced ANTS-Syn based voxel-based analysis method [Schwarz et al., 2014] was also conducted. This method may be less prone to type-I errors than TBSS by utilising single-participant templates and ANTS registration with a white-matter mask rather than a skeletonisation procedure [Schwarz et al., 2014]. This was conducted in accordance with the procedures and parameters detailed by Schwarz et al [Schwarz et al., 2014].

### Results

After correction for multiple comparisons, the ANTS-based analysis revealed only a 4-voxel cluster in the external capsule bordering the putamen. For the TBSS analysis, no voxels showed significant change (p<0.05) after controlling for multiple comparisons. In light of our TMS and tractography findings, we reduced the multiple-comparison threshold of the TBSS analysis to p<0.15 FWE to search for any patterns of change. In this exploratory analysis, a large number of voxels reached threshold in the frontal lobes bilaterally, throughout the presumed right corticomotor tracts, the right and left thalamus, and some areas of the corpus callosum (Supplementary Figure 1).

**Supplementary Figure 1.**
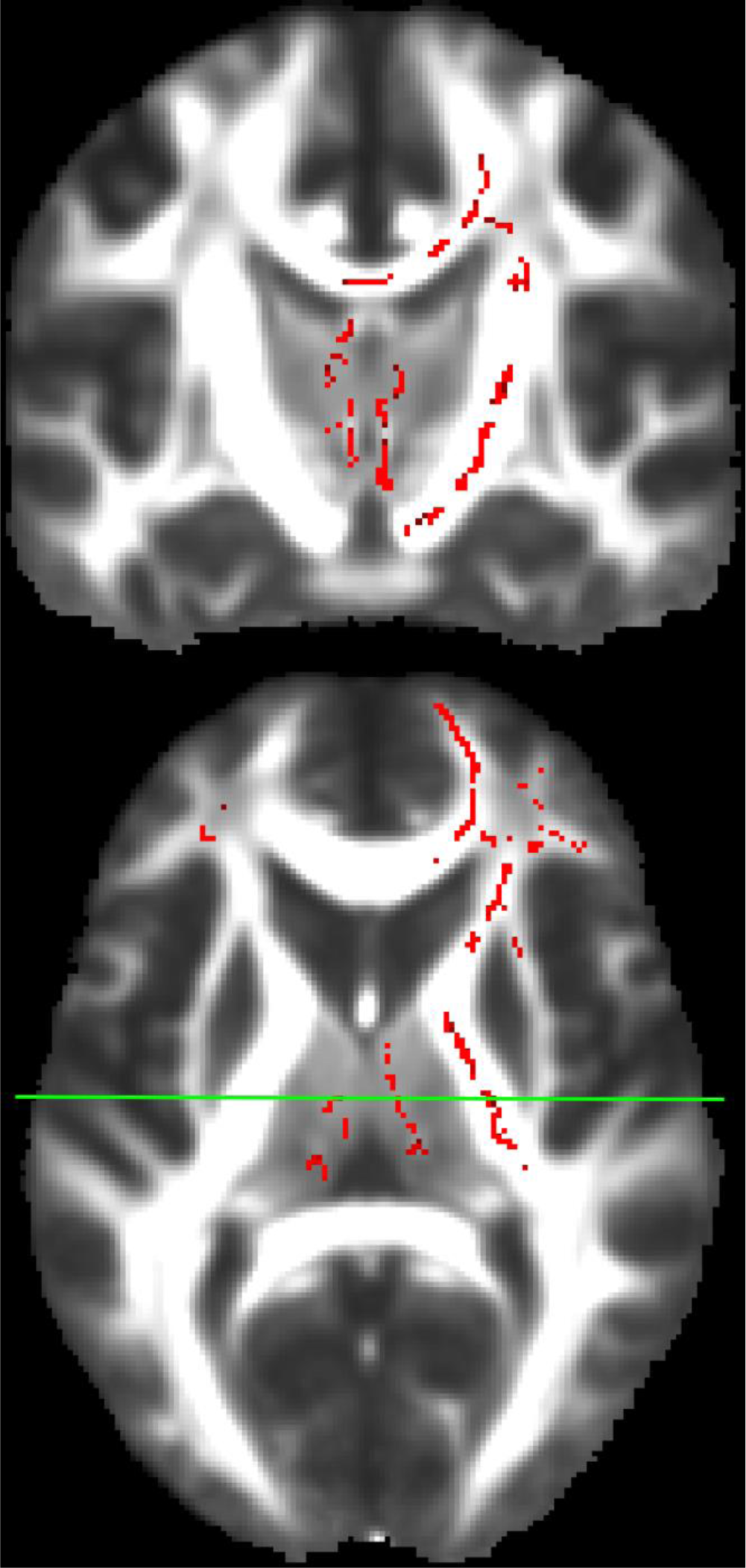
Training-related changes in white matter. ‘Significant’ voxels in the exploratory TBSS analysis (Red) in which the statistical threshold was relaxed to p<0.15 FWE, overlaid with the mean FA image. Voxels consistent with the locations of the right corticomotor tracts, corpus callosum connecting the sensorimotor cortices, bilateral thalamus, and both frontal lobes exceeded this threshold. The top image shows a coronal slice at the level of the line on bottom (axial) image. Right of the image is right of the brain. Voxels did not exceed this threshold in the corticomotor tracts of the ‘untrained’ hemisphere.

## References

Alonso-Ortiz E, Levesque IR, Pike GB (2015): MRI-based myelin water imaging: A technical review. Magn Reson Med 73:70–81. http://www.ncbi.nlm.nih.gov/pubmed/24604728.

Avants BB, Epstein CL, Grossman M, Gee JC (2008): Symmetric diffeomorphic image registration with cross-correlation: evaluating automated labeling of elderly and neurodegenerative brain. Med Image Anal 12:26–41. http://www.ncbi.nlm.nih.gov/pubmed/17659998.

Bach M, Laun FB, Leemans A, Tax CMW, Biessels GJ, Stieltjes B, Maier-Hein KH (2014): Methodological considerations on tract-based spatial statistics (TBSS). Neuroimage 100:358–69. http://dx.doi.org/10.1016/j.neuroimage.2014.06.021.

Baek SO, Jang SH, Lee E, Kim S, Hah JO, Park YH, Lee JM, Son SM (2013): CST recovery in pediatric hemiplegic patients: Diffusion tensor tractography study. Neurosci Lett 557 Pt B:79–83. http://www.ncbi.nlm.nih.gov/pubmed/24176879.

Borich MR, Brodie SM, Gray WA, Ionta S, Boyd LA (2015): Understanding the role of the primary somatosensory cortex: Opportunities for rehabilitation. Neuropsychologia 79:246–55. http://dx.doi.org/10.1016/j.neuropsychologia.2015.07.007.

Chang Y (2014): Reorganization and plastic changes of the human brain associated with skill learning and expertise. Front Hum Neurosci 8:35. http://journal.frontiersin.org/article/10.3389/fnhum.2014.00035/abstract.

Chevrier AD, Noseworthy MD, Schachar R (2007): Dissociation of response inhibition and performance monitoring in the stop signal task using event-related fMRI. Hum Brain Mapp 28:1347–1358.

Dang MT, Yokoi F, Yin HH, Lovinger DM, Wang Y, Li Y (2006): Disrupted motor learning and long-term synaptic plasticity in mice lacking NMDAR1 in the striatum. Proc Natl Acad Sci U S A 103:15254–9. http://www.ncbi.nlm.nih.gov/pubmed/17015831.

Dayan E, Cohen LG (2011): Neuroplasticity subserving motor skill learning. Neuron 72:443–54. http://www.pubmedcentral.nih.gov/articlerender.fcgi?artid=3217208&tool=pmcentrez&rendertype=abstract.

Demerens C, Stankoff B, Logak M, Anglade P, Allinquant B, Couraud F, Zalc B, Lubetzki C (1996): Induction of myelination in the central nervous system by electrical activity. Proc Natl Acad Sci U S A 93:9887–92. http://www.pubmedcentral.nih.gov/articlerender.fcgi?artid=38524&tool=pmcentrez&rendertype=abstract.

Hicks RE, Gualtieri CT, Schroeder SR (1983): Cognitive and Motor Components of Bilateral Transfer. Am J Psychol 96:223. http://www.jstor.org/stable/1422813?origin=crossref.

Ishibashi T, Dakin K a, Stevens B, Lee PR, Kozlov S V, Stewart CL, Fields RD (2006): Astrocytes promote myelination in response to electrical impulses. Neuron 49:823–32. http://www.pubmedcentral.nih.gov/articlerender.fcgi?artid=1474838&tool=pmcentrez&rendertype=abstract.

Jones DK, Symms MR, Cercignani M, Howard RJ (2005): The effect of filter size on VBM analyses of DT-MRI data. Neuroimage 26:546–54. http://www.ncbi.nlm.nih.gov/pubmed/15907311.

Karni A, Meyer G, Jezzard P, Adams MM, Turner R, Ungerleider LG (1995): Functional MRI evidence for adult motor cortex plasticity during motor skill learning. Nature 377:155–8. http://lbcnimh.nih.gov/Ungerleider/Publications/Karni_et_al_Nature_1995.pdf.

Kübler A, Dixon V, Garavan H (2006): Automaticity and reestablishment of executive control-an fMRI study. J Cogn Neurosci 18:1331–42. http://www.ncbi.nlm.nih.gov/pubmed/16859418.

Lu MT, Preston JB, Strick PL (1994): Interconnections between the prefrontal cortex and the premotor areas in the frontal lobe. J Comp Neurol 341:375–92. http://www.ncbi.nlm.nih.gov/pubmed/7515081.

Madhyastha T, Mérillat S, Hirsiger S, Bezzola L, Liem F, Grabowski T, Jäncke L (2014): Longitudinal reliability of tract-based spatial statistics in diffusion tensor imaging. Hum Brain Mapp 35:4544–55. http://www.ncbi.nlm.nih.gov/pubmed/24700773.

Malcom L (2015): Neuroplasticity: how the brain can heal itself. ABC Radio National, April 21. http://www.abc.net.au/radionational/programs/allinthemind/neuroplasticity-and-how-the-brain-can-heal-itself/6406736.

Manjon J V, Coupe P (2016): volBrain: an online MRI brain volumetry system. Article. Front Neuroinform 10. http://www.frontiersin.org/neuroinformatics/10.3389/fninf.2016.00030/abstract.

Mannella F, Gurney K, Baldassarre G (2013): The nucleus accumbens as a nexus between values and goals in goal-directed behavior: a review and a new hypothesis. Front Behav Neurosci 7:135. http://www.ncbi.nlm.nih.gov/pubmed/24167476.

Matyas F, Sreenivasan V, Marbach F, Wacongne C, Barsy B, Mateo C, Aronoff R, Petersen CCH (2010): Motor Control by Sensory Cortex. Science (80) 330:1240–1243. http://www.sciencemag.org/cgi/doi/10.1126/science.1195797.

Modat M, Ridgway GR, Taylor ZA, Lehmann M, Barnes J, Hawkes DJ, Fox NC, Ourselin S (2010): Fast free-form deformation using graphics processing units. Comput Methods Programs Biomed 98:278–84. http://www.sciencedirect.com/science/article/pii/S0169260709002533.

Næss-Schmidt E, Tietze A, Blicher JU, Petersen M, Mikkelsen IK, Coupé P, Manjón J V., Eskildsen SF (2016): Automatic thalamus and hippocampus segmentation from MP2RAGE: comparison of publicly available methods and implications for DTI quantification. Int J Comput Assist Radiol Surg (In Press). http://link.springer.com/10.1007/s11548-016-1433-0.

Palmer HS, Håberg a K, Fimland MS, Solstad GM, Moe Iversen V, Hoff J, Helgerud J, Eikenes L (2013): Structural brain changes after 4 wk of unilateral strength training of the lower limb. J Appl Physiol 115:167–75. http://www.ncbi.nlm.nih.gov/pubmed/23493358.

Pannek K, Boyd RN, Fiori S, Guzzetta A, Rose SE (2014): Assessment of the structural brain network reveals altered connectivity in children with unilateral cerebral palsy due to periventricular white matter lesions. NeuroImage Clin 5:84–92. http://www.pubmedcentral.nih.gov/articlerender.fcgi?artid=4081979&tool=pmcentrez&rendertype=abstract.

Pannek K, Guzzetta A, Colditz PB, Rose SE (2012): Diffusion MRI of the neonate brain: acquisition, processing and analysis techniques. Pediatr Radiol 42:1169–82. http://www.ncbi.nlm.nih.gov/pubmed/22903761.

Penfield W, Boldrey E (1937): Somatic Motor and Sensory Representation in the Cerebral Cortex of Man as Studies by Electrical Stiumlation. Brain 60:389–443. http://brain.oxfordjournals.org/cgi/doi/10.1093/brain/60.4.389.

R Core Team (2015): R: A Language and Environment for Statistical Computing. Vienna, Austria: R Foundation for Statistical Computing. https://www.r-project.org.

Reid LB, Boyd RN, Cunnington R, Rose SE (2016a): Interpreting Intervention Induced Neuroplasticity with fMRI: The Case for Multimodal Imaging Strategies. Neural Plast 2016:1–13. http://www.hindawi.com/journals/np/2016/2643491/.

Reid LB, Cunnington R, Boyd RN, Rose SE (2016b): Surface-Based fMRI-Driven Diffusion Tractography in the Presence of Significant Brain Pathology: A Study Linking Structure and Function in Cerebral Palsy. Ed. Pew-Thian Yap. PLoS One 11:e0159540. http://dx.plos.org/10.1371/journal.pone.0159540.

Sahama I, Sinclair K, Fiori S, Doecke J, Pannek K, Reid L, Lavin M, Rose S (2015): Motor pathway degeneration in young ataxia telangiectasia patients: A diffusion tractography study. NeuroImage Clin 9:206–215. http://linkinghub.elsevier.com/retrieve/pii/S221315821500145X.

Sale M V., Reid LB, Cocchi L, Pagnozzi AM, Rose SE, Mattingley JB Structural and functional brain changes following four weeks of unimanual motor training: evidence from behaviour, neural stimulation, cortical thickness and functional MRI. Hum Brain Mapp VOLUME:PAGES.

Savas JN, Toyama BH, Xu T, Yates JR, Hetzer MW (2012): Extremely long-lived nuclear pore proteins in the rat brain. Science 335:942. http://www.sciencemag.org/content/335/6071/942.abstract.

Scholz J, Klein MC, Behrens TEJ, Johansen-Berg H (2009): Training induces changes in white-matter architecture. Nat Neurosci 12:1370–1371. http://www.pubmedcentral.nih.gov/articlerender.fcgi?artid=2770457&tool=pmcentrez&rendertype=abstract.

Schulder M, Grabow C (2011): The woman who changed her brain. CNN, September. http://cnnradio.cnn.com/2012/09/07/the-woman-who-changed-her-brain/.

Schwarz CG, Reid RI, Gunter JL, Senjem ML, Przybelski SA, Zuk SM, Whitwell JL, Vemuri P, Josephs KA, Kantarci K, Thompson PM, Petersen RC, Jack CR, Alzheimer’s Disease Neuroimaging Initiative (2014): Improved DTI registration allows voxel-based analysis that outperforms tract-based spatial statistics. Neuroimage 94:65–78. http://www.ncbi.nlm.nih.gov/pubmed/24650605.

Siegel JS, Power JD, Dubis JW, Vogel AC, Church J a, Schlaggar BL, Petersen SE (2014): Statistical improvements in functional magnetic resonance imaging analyses produced by censoring high-motion data points. Hum Brain Mapp 35:1981–96. http://www.ncbi.nlm.nih.gov/pubmed/23861343.

Smith RE, Tournier J-D, Calamante F, Connelly A (2015): SIFT2: Enabling dense quantitative assessment of brain white matter connectivity using streamlines tractography. Neuroimage 119:338–51. http://linkinghub.elsevier.com/retrieve/pii/S1053811915005972.

Smith SM, Jenkinson M, Johansen-Berg H, Rueckert D, Nichols TE, Mackay CE, Watkins KE, Ciccarelli O, Cader MZ, Matthews PM, Behrens TEJ (2006): Tract-based spatial statistics: voxelwise analysis of multi-subject diffusion data. Neuroimage 31:1487–505. http://www.ncbi.nlm.nih.gov/pubmed/16624579.

Smith SM, Nichols TE (2009): Threshold-free cluster enhancement: addressing problems of smoothing, threshold dependence and localisation in cluster inference. Neuroimage 44:83–98. http://dx.doi.org/10.1016/j.neuroimage.2008.03.061.

Storr W (2015): The brain’s miracle superpowers of self improvement. BBC Future, November. http://www.bbc.com/future/story/20151123-the-brains-miracle-superpowers-of-self-improvement.

Taubert M, Draganski B, Anwander A, Müller K, Horstmann A, Villringer A, Ragert P (2010): Dynamic properties of human brain structure: learning-related changes in cortical areas and associated fiber connections. J Neurosci 30:11670–7. http://www.ncbi.nlm.nih.gov/entrez/query.fcgi?cmd=Retrieve&db=PubMed&dopt=Citation&list_uids=20810887.

Teixeira L a (2000): Timing and force components in bilateral transfer of learning. Brain Cogn 44:455–69. http://www.ncbi.nlm.nih.gov/pubmed/11104537.

Thomas AG, Marrett S, Saad ZS, Ruff DA, Martin A, Bandettini PA (2009): Functional but not structural changes associated with learning: An exploration of longitudinal Voxel-Based Morphometry (VBM). Neuroimage 48:117–125. http://dx.doi.org/10.1016/j.neuroimage.2009.05.097.

Tournier J-D, Calamante F, Connelly A (2012): MRtrix: Diffusion tractography in crossing fiber regions. Int J Imaging Syst Technol 22:53–66. http://doi.wiley.com/10.1002/ima.22005.

Yeung MSY, Zdunek S, Bergmann O, Bernard S, Salehpour M, Alkass K, Perl S, Tisdale J, Possnert G, Brundin L, Druid H, Frisén J (2014): Dynamics of oligodendrocyte generation and myelination in the human brain. Cell 159:766–74. http://www.ncbi.nlm.nih.gov/pubmed/25417154.

Zatorre RJ, Fields RD, Johansen-Berg H (2012): Plasticity in gray and white: neuroimaging changes in brain structure during learning. Nat Neurosci 15:528–36. http://www.pubmedcentral.nih.gov/articlerender.fcgi?artid=3660656&tool=pmcentrez&rendertype=abstract.

Zhang H, Schneider T, Wheeler-Kingshott C a., Alexander DC (2012): NODDI: Practical in vivo neurite orientation dispersion and density imaging of the human brain. Neuroimage 61:1000–1016. http://dx.doi.org/10.1016/j.neuroimage.2012.03.072.

